## Supplementary Information for "Functionally analogous body- and animacy-responsive areas in the dog (*Canis familiaris*) and human occipito-temporal lobe"

[1]Social, Cognitive and Affective Neuroscience (SCAN) Unit, Department of Cognition, Emotion, and Methods in Psychology, Faculty of Psychology, University of Vienna, Vienna, Austria, [2]Department of Cognitive Biology, Faculty of Life Sciences, University of Vienna, Vienna, Austria, [3]Comparative Cognition, Messerli Research Institute, University of Veterinary Medicine Vienna, Medical University of Vienna and University of Vienna, Vienna, Austria, [4]Vienna Cognitive Science Hub, University of Vienna, Austria

[+]Shared senior authors with equal contributions.

### Validation of threshold to define individual functional regions-of-interest

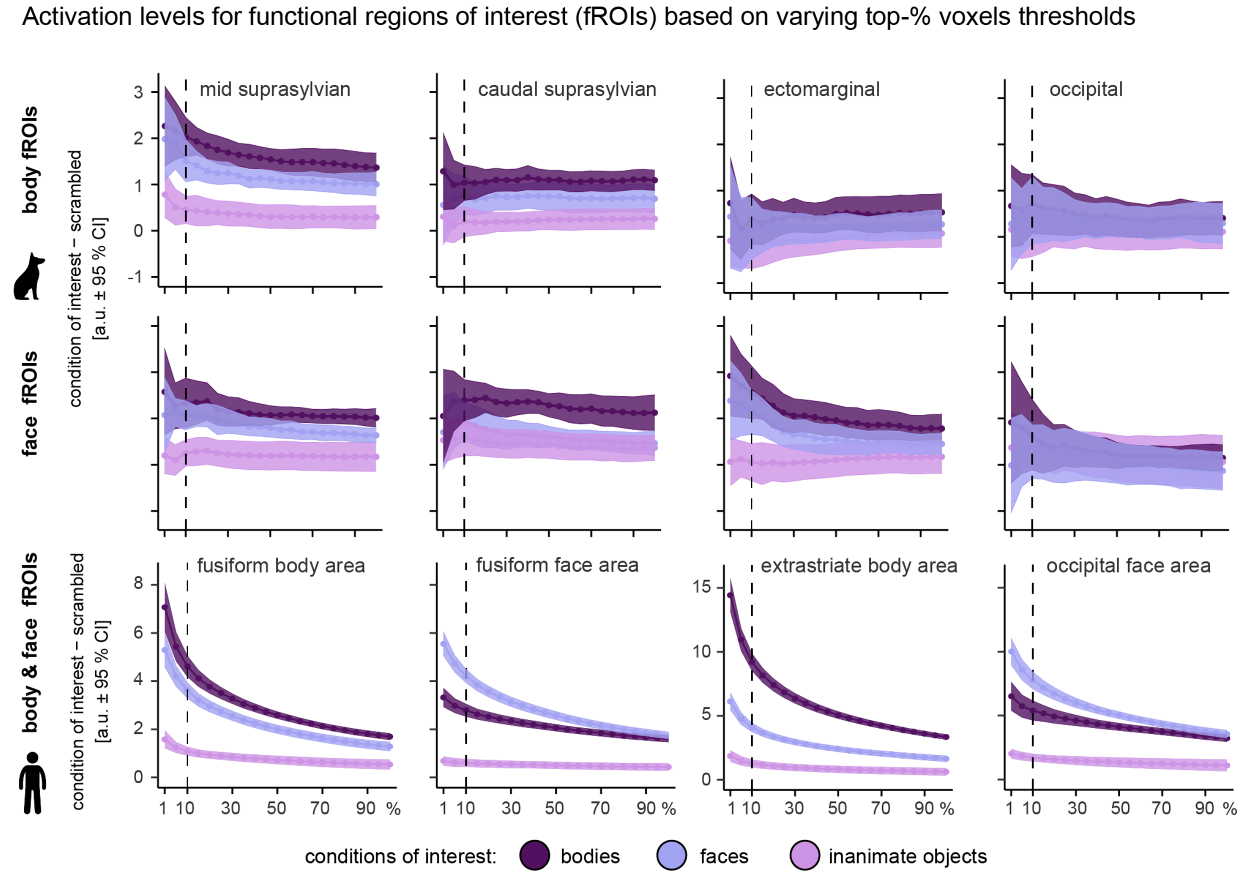


**Supplementary Figure S1.** *Parameter estimates for faces, bodies, and inanimate objects (all contrasts against scrambled controls) retrieved from individual functional regions of interest (fROIs) defined based on top-% most active voxels for faces or bodies > inanimate objects (run 1) ranging from 1% to 100% in steps of 5%.* Smaller percentages would have been less sensitive for dog fROIs and increased fROI sizes would have been less sensitive in the human fusiform and occipital face area as illustrated by the overlapping 95% confidence intervals (CIs). Points represent the mean. a.u., arbitrary units. Dashed line represents the a priori selected 10% threshold used for the main analysis that we set out to validate.

### Control analysis matching the analysis approach used for the dog neuroimaging data

**Supplementary Note 1**

Since anatomical search space definitions and sample sizes differed between the two species, we conducted a control analysis with the human data using anatomical masks instead of parcels (i.e., matching the analysis in the dogs) and performed the analysis in 1000 randomly drawn sub-samples of $n$ = 15 participants (i.e., the dog sample size). We selected the calcarine sulcus including the surrounding cortex, and the fusiform gyrus as anatomical masks to test both primary and higher-order visual regions [1] similar to the dog analysis. Analysis with the full sample of $N$ = 40 human participants revealed no difference in activation levels between faces, bodies, and inanimate objects in the calcarine body fROI and negative activation levels for all three categories compared to baseline (i.e., scrambled control condition) in the calcarine face fROI; differences between faces and bodies in the calcarine fROIs were both non-significant in more than 5% of the sub-samples (see **Supplementary Figures S2-S3**). Results for the fusiform face fROI revealed greater activation levels for faces compared to bodies and inanimate objects in the full sample and a significant difference between faces and bodies in 100% of the sub-sample analyses. The fusiform body fROI also showed greater activation levels for bodies compared to faces and inanimate objects in the full sample and in 97.5% of the sub-samples. Thus, the observed differences between dogs and humans (i.e., face-sensitivity) are also present when accounting for sample size and search space differences (see **Supplementary Figures S2-S3**).

Anatomical search spaces: body functional regions-of-interest (fROIs)

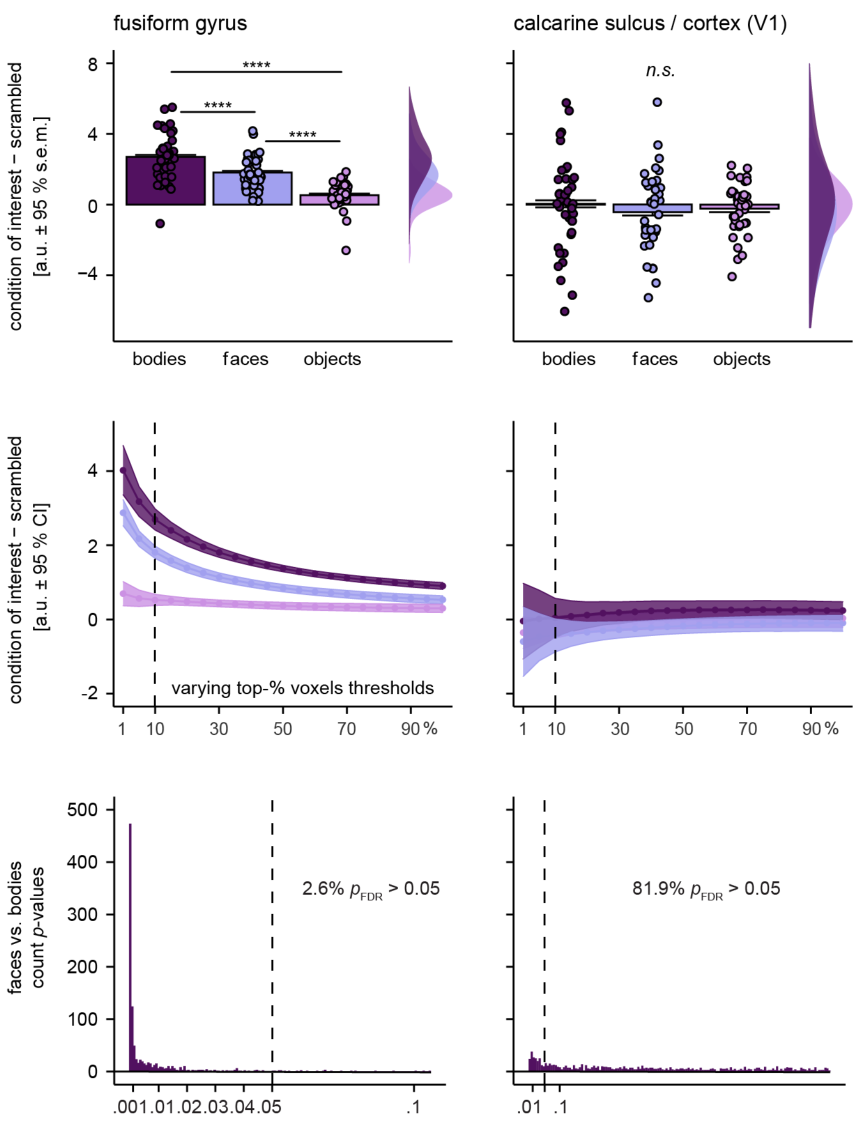


**Supplementary Figure S2.** *Identical functional regions-of-interest (fROI) analysis approach as in dogs revealing a body-sensitive region in the human fusiform gyrus but no category-sensitivity in the primary visual cortex*. Using anatomical masks instead of parcels to define individual body fROIs revealed evidence for body-sensitivity in the human fusiform gyrus ($F(1.90,74.25) = 115.31$, $p_{FDR} < .0001$, $\eta^2_g = .443$) but not the calcarine cortex ($F(1.41,77.79) = 1.43$, $p_{FDR} < .245$, $\eta^2_g = .008$) in the full sample of $N = 40$ human participants (first row). Second row: Visual inspection of parameter estimates for faces, bodies and inanimate objects (all > scrambled images) confirms top-10% active voxels as sufficient fROI defining threshold for all fROIs. Running the analysis with 1000 randomly drawn sub-samples of $n = 15$ human participants (i.e., equivalent to dog sample size) confirmed again greater activation for bodies compared to faces in 97.4% of the cases (i.e., less than 5% nonsignificant comparisons) in the fusiform body fROI and no significant difference between bodies and faces in calcarine fROI in 81.9% of the cases. Planned comparisons were false discovery rate (FDR) corrected to control for multiple comparisons. ****$p_{FDR} < .0001$, error bars represent the standard error of the mean (left panel) or the 95% confidence interval (CI) of the mean (s.e.m.) in the middle panel; n.s., not significant; a.u., arbitrary units; V1, primary visual cortex

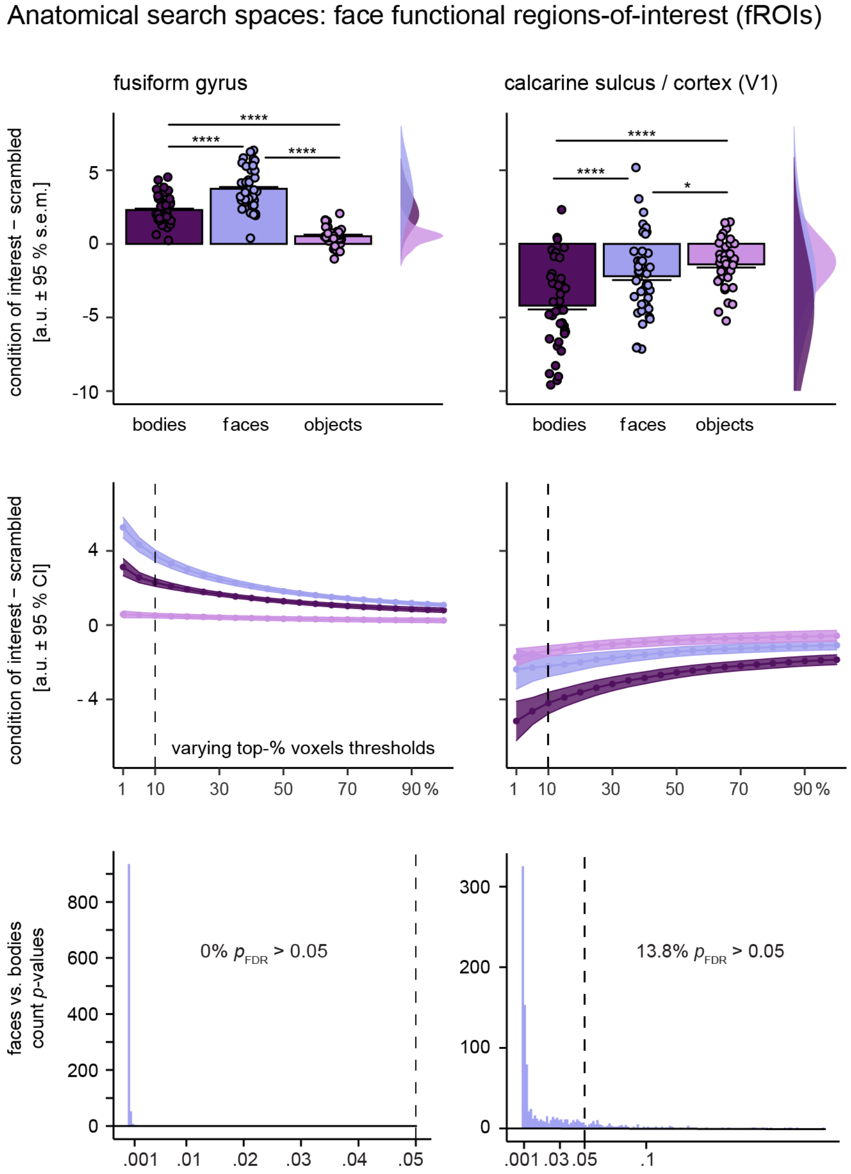


**Supplementary Figure S3.** *Identical functional regions-of-interest (fROI) analysis approach as in dogs revealing a face-sensitive region in the human fusiform gyrus but not in the primary visual cortex*. Using anatomical masks instead of parcels to define individual face fROIs, revealed evidence for face-sensitivity in the human fusiform gyrus ($F$(1.82,70.86) = 214.45, $p_{FDR}$ < .0001, $\eta^2_g$.= .622) in the full sample of $N$ = 40 human participants (first row). In the calcarine cortex, activation levels between faces and bodies were significantly different ($F$(1.94,75.68) = 34.01, $p_{FDR}$ < .0001, $\eta^2_g$.= .187) but activation levels for faces, bodies, and inanimate objects were negative indicating less for all categories compared to scrambled images. Second row: Visual inspection of parameter estimates for faces, bodies, and inanimate objects (all > scrambled images) confirms top-10% active voxels as sufficient fROI defining threshold for all fROIs. Running the analysis with 1000 randomly drawn sub-samples of $n$ = 15 human participants (i.e., equivalent to dog sample size) confirmed again greater activation for faces compared to bodies in 100% of the cases in the fusiform face fROI. In the calcarine cortex differences between faces and bodies were not significant in 13.8%. Planned comparisons were false discovery rate (FDR) corrected to control for multiple comparisons. *$p_{FDR}$ < .05, ****$p_{FDR}$ < .0001, error bars represent the standard error of the mean (left panel) or the 95% confidence interval (CI) of the mean (s.e.m.) in the middle panel; a.u., arbitrary units; V1, primary visual cortex.

### Exploratory analysis of low-level visual properties of the stimulus set

### Supplementary Note 2

In order to control for low-level visual differences, the stimulus set had been controlled for size, spatial extent, and luminance (see **Stimulus Material** for detailed information). We did not control for further low-level visual properties to preserve ecological validity (i.e., dog fur and human clothing do differ in hue in everyday life). However, to test if the observed effects might have been driven by low-level visual properties, we conducted exploratory analyses comparing hue, saturation and contrast measures between face, body, and inanimate object stimuli and between dog and human face and body stimuli (i.e., mirroring the fROI analysis). Results revealed no differences in contrast between stimuli categories but expected differences in hue and saturation (see **Figure S4**, **Supplementary Tables S7-S9**): hue differed between bodies and inanimate objects as well as between faces and inanimate objects, but not between faces and bodies. Comparing dog and human stimuli, we found a significant difference in hue between dog and human bodies. Faces and inanimate objects did not differ in saturation, but body stimuli had a lower saturation compared to faces and inanimate objects; and dog faces had a lower saturation compared to human faces. Overall, these findings indicate that the observed differences in the dog and human fROI analysis do not reflect low-level visual differences. For example, if dog fROIs were driven by differences in hue and saturation we should have also seen differences in activation levels in response to human and dog bodies (i.e., hue) or dog and human faces (i.e., saturation).

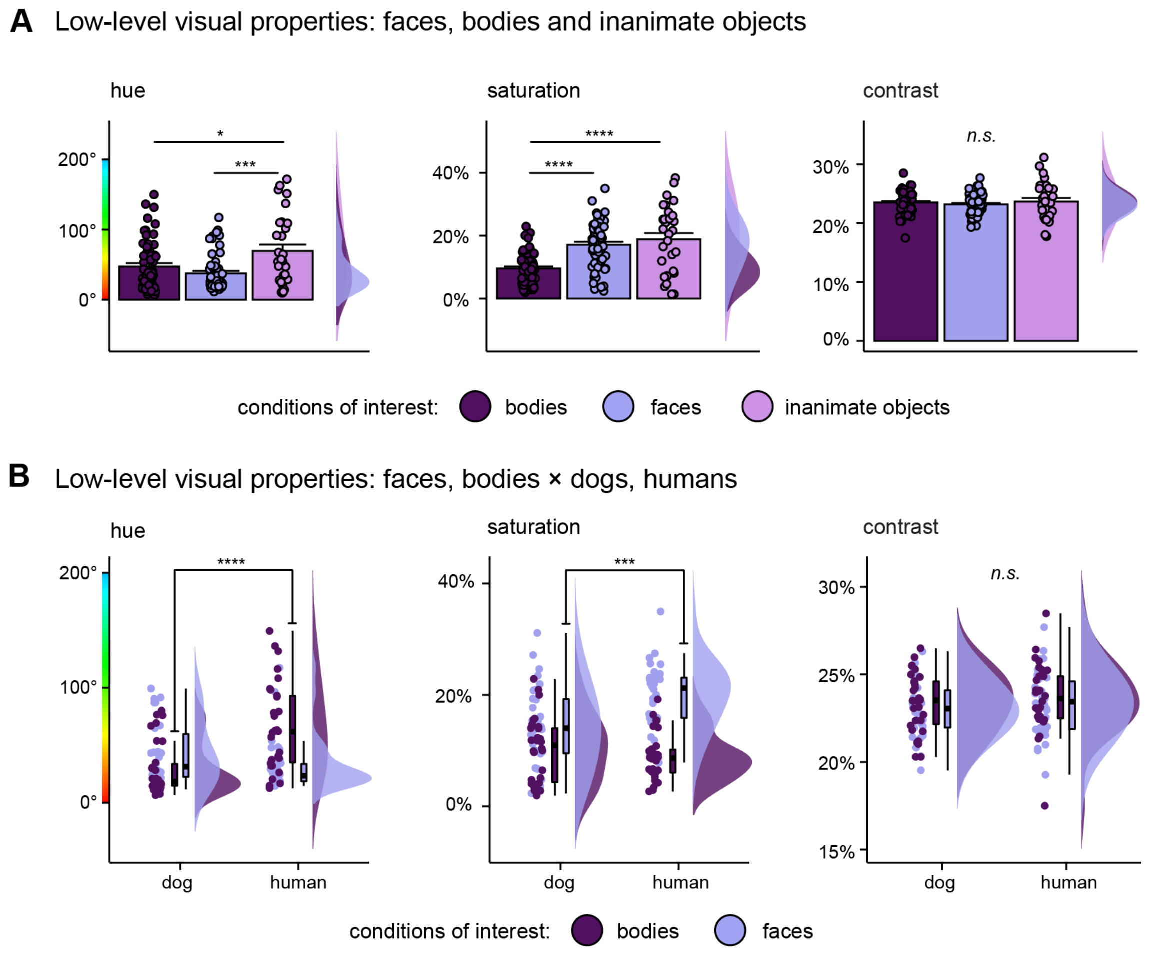


**Supplementary Figure S4.** *Exploratory analysis shows low-level visual differences between conditions for hue and saturation, but not for contrast.* Hue represents degrees (°) on a colour wheel (range: 0° to 360°), saturation and contrast are measured in percentages (range: 0% to 100%). (**A**) Hue of face and body stimuli each differ from inanimate objects, (**B**) adding species as factor further showed a difference between dog and human body stimuli. (**A**) Body stimuli have significantly lower saturation than face or inanimate object stimuli. The latter do not differ in saturation, (**B**) but within the face stimuli, saturation levels are significantly higher for human compared to dog faces. (**A B**) We did not find any differences between stimuli categories for contrast. Detailed information can be found in **Supplementary Tables S7-S8**. The stimulus set has been controlled for luminance, spatial extent (= ratio image / background) and size (see section **Stimulus material** for details). Post-hoc comparisons were false discovery rate (FDR) corrected to control for multiple comparisons *$p$ < .05, ***$p$ < .001, ****$p$ < .0001.

**Supplementary Table S1.** Task-related activation: localizer data set (task run 1)

| Contrast & brain region | Coordinates | | | *z*-value | cluster size |
|---|---|---|---|---|---|
| | x | y | z | | |
| **All conditions > implicit visual baseline** | | | | | |
| **Dog participants** | | | | | |
| R mid suprasylvian gyrus | -16 | -22 | 14 | 4.56 | 416 |
| L caudal suprasylvian gyrus | -22 | -26 | 0 | 4.39 | 44 |
| R caudal suprasylvian gyrus | 22 | -24 | 0 | 4.04 | 45 |
| **Human participants** | | | | | |
| R middle occipital gyrus | 48 | -70 | -8 | Inf | 40800 |
| R middle frontal gyrus | 54 | 12 | 40 | 5.41 | 1360 |
| L middle temporal gyrus | -64 | -8 | -16 | 5.31 | 73 |
| L superior frontal gyrus | -4 | 14 | 56 | 4.72 | 128 |
| R superior frontal gyrus | 4 | 14 | -12 | 4.69 | 148 |
| L superior frontal gyrus | -10 | 62 | 48 | 4.57 | 78 |

*Note.* Effects were tested for significance with a cluster defining threshold of $p$ <.005/.001 (dogs/humans) and a cluster probability threshold of $p$ < .05 FWE corrected for multiple comparisons. We report the first local maximum within each cluster of each sample *t*-test. Coordinates for the dog data represent the location of the peak voxels referring to the canine breed-averaged template space[2]; the template along with another detailed dog anatomical atlas[3] served to determine anatomical nomenclature for the dog data. Human data coordinates refer to the stereotactic Montreal Neurological Institute (MNI) space, we obtained anatomical nomenclature from the Laboratory for Neuro Imaging (LONI) Brain Atlas[4], LBPA40, http://www.loni.usc.edu/atlases/). Data of both samples is presented in **Figure 1C**. L, left; R, right.

**Supplementary Table S2.** fROI sizes with top-10% active voxels cut-off

| | left | | right | |
|---|---|---|---|---|
| Sample, search space and fROI | mean | *SD* | mean | *SD* |
| **Dog participants: body fROIs** | | | | |
| Mid suprasylvian gyrus | 8.33 | 2.97 | 11.13 | 3.7 |
| Caudal suprasylvian gyrus | 10.73 | 2.76 | 10 | 4.49 |
| Ectomarginal gyrus | 9.47 | 4.32 | 9.2 | 3.76 |
| Marginal gyrus | 8.93 | 3.45 | 8.87 | 2.77 |
| Splenial gyrus | 12.73 | 5.35 | 8.8 | 4.23 |
| Occipital gyrus | 4.6 | 2.03 | 5.52 | 3.09 |
| **Dog participants: face fROIs** | | | | |
| Mid suprasylvian gyrus | 10.6 | 3.36 | 9.07 | 3.61 |
| Caudal suprasylvian gyrus | 8.4 | 2.87 | 7.2 | 3.34 |
| Ectomarginal gyrus | 11.47 | 4.73 | 9.07 | 3.56 |
| Marginal gyrus | 9.8 | 4.3 | 12.33 | 3.37 |
| Splenial gyrus | 14.27 | 5.18 | 12.4 | 4.64 |
| Occipital gyrus | 5.93 | 2.58 | 7.73 | 2.63 |
| **Human participants: body fROIs** | | | | |
| Extrastriate body area | 168.25 | 30.04 | 204.8 | 29.09 |
| Fusiform body area | 70.85 | 29.24 | 84 | 25.4 |
| Anatomical mask: fusiform body area | 71.62 | 27.09 | 79.35 | 28.66 |
| **Human participants: face fROIs** | | | | |
| Occipital face area | 14.72 | 15.83 | 48.5 | 15.97 |
| Fusiform face area | 28.73 | 10.92 | 64 | 16.09 |
| Anatomical mask: fusiform face area | 69 | 30.82 | 72.17 | 30.95 |

*Note*. Summary statistics for individual face and body functional regions-of-interest (fROIs) split for right and left hemisphere.*SD*, standard deviation.

**Supplementary Table S3.** Results from one-way[a] repeated measures analyses of variance (ANOVAs)

| Search space (predictor: image category) | $df_{\text{Num}}$ | $df_{\text{Den}}$ | F | $p$ | $p_{\text{FDR}}$ | $\eta^2_g$ |
|---|---|---|---|---|---|---|
| **Dog participants** | | | | | | |
| **Individual body fROIs** | | | | | | |
| Mid suprasylvian gyrus | 1.77 | 24.78 | 23.83 | **< .0001** | **< .0001** | .63 |
| Caudal suprasylvian gyrus | 1.60 | 22.47 | 7.16 | **.006** | **.018** | .34 |
| Ectomarginal gyrus | 2 | 27.99 | 1.23 | .308 | .37 | .081 |
| Marginal gyrus | 1.85 | 25.96 | 1.19 | .317 | .317 | .078 |
| Splenial gyrus | 1.98 | 27.79 | 2.06 | .146 | .292 | .129 |
| Occipital gyrus | 1.89 | 26.45 | 1.56 | .230 | .345 | .10 |
| **Individual face fROIs** | | | | | | |
| Mid suprasylvian gyrus | 1.77 | 24.75 | 8.19 | **.003** | **.009** | .369 |
| Caudal suprasylvian gyrus | 1.93 | 26.96 | 4.56 | **.021** | **.042** | .246 |
| Ectomarginal gyrus | 1.77 | 24.81 | 9.41 | **.001** | **.006** | .402 |
| Marginal gyrus | 1.85 | 25.87 | 2.60 | .097 | .146 | .157 |
| Splenial gyrus | 1.42 | 19.82 | .57 | .518 | .622 | .039 |
| Occipital gyrus | 1.97 | 27.52 | .40 | .671 | .671 | .028 |
| **Human participants** | | | | | | |
| **Individual body fROIs** | | | | | | |
| Extrastriate body area | 1.44 | 56.19 | 264 | **< .0001** | **< .0001** | .871 |
| Fusiform body area | 1.92 | 75.02 | 129.9 | **< .0001** | **< .0001** | .769 |
| **Individual face fROIs** | | | | | | |
| Occipital face area | 1.87 | 72.77 | 108.63 | **< .0001** | **< .0001** | .736 |
| Fusiform face area | 1.94 | 75.85 | 206.35 | **< .0001** | **< .0001** | .841 |

*Note.* [a]Independent variable: *image category* (levels: bodies, faces, inanimate objects; all over scrambled control), dependent variable: activation levels. For each sample (i.e., dogs and humans) *p*-values for group comparisons investigating the same research question (i.e., body-preference for bodies) are FDR (false discovery rate) corrected, uncorrected *p*-values are also reported. *P*-values < .05 are in bold. Post-hoc comparison results are presented in **Figure 2B**. $df_{Num}$, degrees of freedom numerator; $df_{Den}$ degrees of freedom denominator; $\eta^2_g$, generalized eta-squared.

**Supplementary Table S4.** Whole-brain univariate analysis: face-, body- and animacy-sensitivity (dog data)

| Contrast & brain region | Coordinates | | | z-value | cluster size |
|---|---|---|---|---|---|
| | x | y | z | | |
| **Bodies > faces, inanimate objects** | | | | | |
| L mid suprasylvian gyrus | -16 | -22 | 13 | 5.63 | 67 |
| R caudal suprasylvian gyrus | 20 | -26 | 4 | 4.48 | 129 |
| L caudal suprasylvian gyrus | -20 | -24 | 2 | 4.33 | 42 |
| **Animate (faces, bodies) > inanimate objects** | | | | | |
| L mid suprasylvian gyrus | -18 | -24 | 16 | 5.35 | 60 |

*Note.* Effects were tested for significance with a cluster defining threshold of $p$ <.005 and a cluster probability threshold of $p$ < .05 FWE corrected for multiple comparisons. We report the first local maximum within each cluster for each post hoc sample $t$-tests, the critical cluster size to determine significance was $k$ = 39. Post-hoc comparisons for faces > bodies, inanimate objects did not survive multiple comparison corrections. Coordinates represent the location of the peak voxels referring to the canine breed-averaged template space[2]; the template along with another detailed dog atlas[3] served to determine anatomical nomenclature. The corresponding activation maps are visualized in **Figure 3A**. L, left; R, right.

**Supplementary Table S5.** Whole-brain univariate analysis: face-, body- and animacy-sensitivity (human data)

| Contrast & brain region | Coordinates | | | *z*-value | cluster size |
|---|---|---|---|---|---|
| | x | y | z | | |
| **Bodies > faces, inanimate objects** | | | | | |
| L middle occipital gyrus | -46 | -70 | 10 | Inf | 20866 |
| R middle frontal gyrus | 30 | -2 | 50 | 6.16 | 1303 |
| R lingual gyrus | 22 | -64 | -4 | 6.05 | 1050 |
| L thalamus | -18 | -30 | 4 | 6.03 | 87 |
| L middle frontal gyrus | -28 | -6 | 52 | 5.9 | 979 |
| **Faces > bodies, inanimate objects** | | | | | |
| R fusiform gyrus | 42 | -50 | -22 | 7.8 | 999 |
| R hippocampus | 18 | -4 | -16 | 7.08 | 864 |
| L inferior occipital gyrus | -38 | -88 | -14 | 6.90 | 454 |
| L hippocampus | -18 | -4 | -16 | 6.44 | 638 |
| R precuneus | 4 | -54 | 32 | 5.99 | 741 |
| **Animate (faces, bodies) > inanimate objects** | | | | | |
| R middle occipital gyrus | 46 | -70 | -4 | Inf | 5144 |
| L middle occipital gyrus | -48 | -80 | -2 | Inf | 4554 |
| R hippocampus | 18 | -4 | -14 | 6.96 | 896 |
| L hippocampus | -18 | -6 | -16 | 6.35 | 651 |
| R superior parietal gyrus | 30 | -48 | 56 | 6.12 | 4189 |

*Note.* Effects were tested for significance with a cluster defining threshold of $p < .001$ and a cluster probability threshold of $p < .05$ FWE corrected for multiple comparisons. We report the first local maximum within each cluster for each post hoc sample *t*-tests limited to the first five cluster with the highest peak *z*-values; the critical cluster size to determine significance was $k = 66$ voxels. Coordinates represent the location of the peak voxels referring to the stereotactic Montreal Neurological Institute (MNI) space. We obtained anatomical nomenclature from the Laboratory for Neuro Imaging (LONI) Brain Atlas[4], LBPA40, http://www.loni.usc.edu/atlases/) and the Automatic Anatomical Labelling atlas (AAL2; Rolls, Joliot and Tzourio-Mazoyer, 2015). Data is presented in **Figure 3B**. L, left; R, right.

**Supplementary Table S6.** Results from 2 × 2 repeated measures analyses of variance (ANOVAs) [a]

| Search space, predictor | $df_{Num}$ | $df_{Den}$ | F | *p* | $p_{FDR}$ | $\eta^2_g$ |
|---|---|---|---|---|---|---|
| **Dog participants** | | | | | | |
| **Individual body fROIs** | | | | | | |
| Mid suprasylvian gyrus | | | | | | |
| Species | 1 | 14 | 1.40 | .257 | .514 | .091 |
| Animate stimulus | 1 | 14 | 7.19 | **.018** | .108 | .339 |
| Species × animate stimulus | 1 | 14 | 1.37 | .261 | .392 | .089 |
| Caudal suprasylvian gyrus | | | | | | |
| Species | 1 | 14 | 4.70 | **.048** | .144 | .251 |
| Animate stimulus | 1 | 14 | 1.08 | .317 | .380 | .071 |
| Species × animate stimulus | 1 | 14 | .03 | .855 | .855 | .002 |
| **Individual face fROIs** | | | | | | |
| Mid suprasylvian gyrus | | | | | | |
| Species | 1 | 14 | .99 | .337 | .433 | .066 |
| Animate stimulus | 1 | 14 | 1.06 | .32 | .48 | .071 |
| Species × animate stimulus | 1 | 14 | 1.81 | .2 | .36 | .114 |
| Caudal suprasylvian gyrus | | | | | | |
| Species | 1 | 14 | 0 | .995 | .995 | < .001 |
| Animate stimulus | 1 | 14 | 2.95 | .108 | .324 | .174 |
| Species × animate stimulus | 1 | 14 | .41 | .535 | .602 | .028 |
| Ectomarginal gyrus | | | | | | |
| Species | 1 | 14 | 3.55 | .08 | .72 | .202 |
| Animate stimulus | 1 | 14 | 1.83 | .198 | .446 | .116 |
| Species × animate stimulus | 1 | 14 | 3.25 | .093 | .419 | .188 |
| **Human participants** | | | | | | |
| **Individual body fROIs** | | | | | | |
| Extrastriate body area | | | | | | |
| Species | 1 | 39 | 3.29 | .077 | .77 | .078 |
| Animate stimulus | 1 | 39 | 219.45 | **< .0001** | **< .0001** | .849 |
| Species × animate stimulus | 1 | 39 | 15.33 | **< .0001** | **< .0001** | .282 |
| Fusiform body area | | | | | | |
| Species | 1 | 39 | .31 | .579 | .695 | .008 |
| Animate stimulus | 1 | 39 | 63.43 | **< .0001** | **< .0001** | .312 |
| Species × animate stimulus | 1 | 39 | 4.41 | **.042** | .063 | .044 |
| **Individual face fROIs** | | | | | | |
| Occipital face area | | | | | | |
| Species | 1 | 39 | .01 | .917 | .917 | < .001 |
| Animate stimulus | 1 | 39 | 69.57 | **< .0001** | **< .0001** | .641 |
| Species × animate stimulus | 1 | 39 | 1.36 | .251 | .301 | .034 |
| Fusiform face area | | | | | | |
| Species | 1 | 39 | 15.77 | **< .001** | **< .001** | .288 |
| Animate stimulus | 1 | 39 | 102.16 | **< .0001** | **< .0001** | .724 |
| Species × animate stimulus | 1 | 39 | 1.44 | .237 | .356 | .036 |

*Note.* [a]Independent variables are *species* (levels: dog, human) and *animate stimulus* (levels: bodies, faces), all over scrambled control; dependent variable: activation levels. For each participant group, *p*-values for group comparisons investigating the same research question (i.e., species-preference) are FDR (false discovery rate) corrected, uncorrected *p*-values are also reported. *P*-values < .05 are in bold. Post-hoc comparison results are presented in **Figure 4**. $df_{Num}$, degrees of freedom numerator; $df_{Den}$ degrees of freedom denominator; $\eta^2_g$, generalized eta-squared.

**Supplementary Table S7.** Results from one-way analyses of variance (ANOVAs)[a]

| Visual property (predictor: image category) | $df_{Num}$ | $df_{Den}$ | F | *p* | $\eta^2_g$ |
|---|---|---|---|---|---|
| **Hue** | 2 | 147 | 7.94 | **< .001** | .098 |
| **Saturation** | 2 | 147 | 22.28 | **< .001** | .233 |
| **Contrast** | 2 | 147 | .58 | .560 | .008 |

*Note.* [a]Independent variable: *image category* (levels: bodies, faces, inanimate objects), dependent variable: measurement of low-level visual property (i.e., hue, saturation, contrast). *P*-values < .05 are displayed in bold. Post-hoc comparison results are presented in **Figure S4A**. $df_{Num}$, degrees of freedom numerator; $df_{Den}$ degrees of freedom denominator; $\eta^2_g$, generalized eta-squared.

**Supplementary Table S8.** Results from two-way analyses of variance (ANOVAs)[a]

| Measurement, predictor | $df_{Num}$ | $df_{Den}$ | F | *p* | $\eta^2_g$ |
|---|---|---|---|---|---|
| **Hue** | | | | | |
| Species | 1 | 116 | 5.45 | **.021** | .045 |
| Animate stimulus | 1 | 116 | 3.56 | .062 | .03 |
| Species × animate stimulus | 1 | 116 | 20.24 | **<.001** | .149 |
| **Saturation** | | | | | |
| Species | 1 | 14 | 4.39 | **.038** | .036 |
| Animate stimulus | 1 | 14 | 48.36 | **<.001** | .294 |
| Species × animate stimulus | 1 | 14 | 10.87 | **<.001** | .086 |
| **Contrast** | | | | | |
| Species | 1 | 14 | .76 | .329 | .007 |
| Animate stimulus | 1 | 14 | .96 | .384 | .008 |
| Species × animate stimulus | 1 | 14 | 0 | .950 | <.001 |

*Note.* [a]Independent variables: *animate stimulus* (levels: bodies, faces), *species* (levels: dogs, humans), dependent variable: measurement of low-level visual property (i.e., hue, saturation, contrast). *P*-values < .05 are displayed in bold. Post-hoc comparison results are presented in **Figure S4B.** $df_{Num}$, degrees of freedom numerator; $df_{Den}$ degrees of freedom denominator; $\eta^2_g$, generalized eta-squared.

**Supplementary Table S9.** Dog data: Pattern similarity during face, body, and object perception

| Contrast & brain region | Coordinates | | | *t*-value | cluster size |
|---|---|---|---|---|---|
| | x | y | z | | |
| Bodies [dog bodies × human bodies] > inanimate objects | | | | | |
| R caudal suprasylvian gyrus | 18 | -24 | 4 | 5.77 | 116 |
| Faces [dog faces × human faces] > inanimate objects | | | | | |
| R marginal gyrus | 5 | -34 | 20 | 7.01 | 245 |
| R olfactory tuberculum | 8 | 8 | -2 | 6.16 | 80 |
| Conspecific (dog) bodies > heterospecific (human) bodies | | | | | |
| R mid ectosylvian gyrus | 16 | -16 | 12 | 5.53 | 62 |
| L caudal suprasylvian gyrus | -16 | -27 | 4 | 5.21 | 99 |
| L piriform lobe | -14 | -3 | -4 | 5.18 | 95 |
| Animate [dog bodies × human bodies × dog faces × human faces] > inanimate stimuli | | | | | |
| R splenial gyrus | 5 | -32 | 7 | 5.38 | 70 |
| R caudal suprasylvian gyrus | 18 | -24 | 2 | 5.25 | 72 |

*Note.* Effects were tested for significance with a cluster defining threshold of $p$ <.005 and a cluster probability threshold of $p < .05$ FWE corrected for multiple comparisons. We report the first local maximum within each cluster for all paired sample *t*-tests. The results from the paired sample *t*-tests comparing faces [dog faces × human faces] vs. bodies [dog bodies × human bodies] and conspecific (dog) faces vs. heterospecific (human) faces along with the reversed contrasts [dog bodies × human bodies] < inanimate objects, [dog faces × human faces] < inanimate objects, conspecific (dog) bodies < heterospecific (human) bodies and [dog bodies × human bodies × dog faces × human faces] < inanimate objects did not survive the statistical threshold. Coordinates represent the location of the peak voxels referring to the canine breed-averaged template space[2]; the template along with another dog atlas[3] served to determine anatomical nomenclature for the dog data. Data is presented in **Figure 5**. L, left; R, right.

**Supplementary Table S10.** Human data: Pattern similarity during face, body, and object perception

| Contrast & brain region | Coordinates | | | *t*-value | cluster size |
|---|---|---|---|---|---|
| | x | y | z | | |
| Bodies [dog bodies × human bodies] > inanimate objects | | | | | |
| R middle occipital gyrus | 56 | -72 | 6 | 14.98 | 14018 |
| L middle occipital gyrus | -54 | -78 | 10 | 12.32 | 1622 |
| L cerebellum | -4 | -70 | -40 | 7.33 | 585 |
| L superior parietal gyrus | -26 | -62 | 68 | 6.65 | 1383 |
| R superior frontal gyrus | 6 | 62 | -4 | 6.17 | 881 |
| L precentral gyrus | -26 | -10 | 50 | 5.26 | 570 |
| Bodies [dog bodies × human bodies] < inanimate objects | | | | | |
| L inferior temporal gyrus | -22 | -48 | -12 | 11.26 | 6678 |
| Faces [dog faces × human faces] > inanimate objects | | | | | |
| R fusiform gyrus | 46 | -42 | -20 | 9.35 | 1088 |
| R middle occipital gyrus | 56 | -72 | 0 | 5-71 | 349 |
| Faces [dog faces × human faces] < inanimate objects | | | | | |
| L cerebellum | -20 | -54 | -18 | 14.20 | 7002 |
| R lingual gyrus | 24 | -56 | -8 | 11.84 | 5198 |
| L precentral gyrus | -46 | 6 | 32 | 4.77 | 353 |
| Faces [dog faces × human faces] > bodies [dog bodies × human bodies] | | | | | |
| R lingual gyrus | 6 | -88 | 0 | 11.35 | 3461 |
| Faces [dog faces × human faces] < bodies [dog bodies × human bodies] | | | | | |
| R middle occipital gyrus | 34 | -72 | 10 | 14.21 | 39057 |
| R middle orbitofrontal gyrus | 24 | 42 | -14 | 5.53 | 1087 |
| R cerebellum | 8 | -72 | -38 | 5.19 | 572 |
| R cingulate gyrus | 2 | 6 | 26 | 4.76 | 267 |
| Conspecific (human) bodies > heterospecific (dog) bodies | | | | | |
| R fusiform gyrus | 44 | -44 | -22 | 5.76 | 1053 |
| L middle occipital gyrus | -54 | -72 | 12 | 4.87 | 267 |
| R middle temporal gyrus | 40 | -54 | 12 | 4.81 | 468 |
| R superior parietal gyrus | 16 | -54 | 42 | 4.60 | 466 |
| Heterospecific (dog) faces > conspecific (human) faces | | | | | |
| L middle occipital gyrus | -16 | -94 | 6 | 7.20 | 8147 |
| Brainstem | 6 | -40 | -26 | 4.64 | 618 |
| Animate [dog bodies × human bodies × dog faces × human faces] > inanimate objects | | | | | |
| R middle occipital gyrus | 56 | -72 | 0 | 9.09 | 2035 |
| L cerebellum | -4 | -70 | -42 | 7.30 | 277 |
| R postcentral gyrus | 36 | -26 | 48 | 5.09 | 403 |
| R superior parietal gyrus | 38 | -44 | 68 | 4.82 | 338 |
| R superior frontal gyrus | 8 | 52 | -8 | 4.77 | 564 |
| Animate [dog bodies × human bodies × dog faces × human faces] < inanimate objects | | | | | |
| L cerebellum | -22 | -50 | -18 | 12.63 | 11878 |

*Note.* Effects were tested for significance with a cluster defining threshold of $p$ <.001 and a cluster probability threshold of $p$ < .05 FWE corrected for multiple comparisons. We report the first local maximum within each cluster for all paired sample *t*-tests. Results from the reversed contrasts conspecific (human) bodies < heterospecific (dog) bodies and heterospecific (dog) faces < conspecific (human) faces did not survive the statistical threshold. Coordinates represent the location of the peak voxels and we obtained anatomical nomenclature from the Laboratory for Neuro Imaging (LONI) Brain Atlas[4], LBPA40, http://www.loni.usc.edu/atlases/). Data is presented in **Figure 5.**

**Supplementary References**


1. Grill-Spector, K. & Malach, R. The Human Visual Cortex. *Annu. Rev. Neurosci.* **27**, 649–677 (2004).
2. Nitzsche, B. *et al.* A stereotaxic breed-averaged, symmetric T2w canine brain atlas including detailed morphological and volumetrical data sets. *Neuroimage* **187**, 93–103 (2019).
3. Czeibert, K., Andics, A., Petneházy, Ö. & Kubinyi, E. A detailed canine brain label map for neuroimaging analysis. *Biol. Futur.* **70**, 112–120 (2019).
4. Shattuck, D. W. *et al.* Construction of a 3D probabilistic atlas of human cortical structures. *Neuroimage* **39**, 1064–1080 (2008).
5. Rolls, E. T., Joliot, M. & Tzourio-Mazoyer, N. Implementation of a new parcellation of the orbitofrontal cortex in the automated anatomical labeling atlas. *Neuroimage* **122**, 1–5 (2015).
